# Spatial transcriptomics reveals selective vulnerability of cardiac neural crest-derived glial cells in human myocardial infarction

**DOI:** 10.64898/2026.09.14.751638

**Authors:** Xuefeng Xia, Yaping Ding, Xiaopeng Zhao, Qian He, Hongru Zhang, Yihuang Gu, Hua Bai, Daoyou Pan

## Abstract

Cardiac neural crest-derived glial cells (CNGs) are recently identified glial cells that support cardiac autonomic innervation and maintain sympathetic-parasympathetic balance. Their fate in myocardial infarction (MI) is unknown. Here, we analyzed 16 human cardiac spatial transcriptomes spanning normal myocardium (n=4), infarct zone (n=5), border zone (inner, n=1; outer, n=3), and remote zone (n=3). CNGs (S100B^+^/GFAP^+^, CD68^−^ spots) were selectively lost along a spatial gradient from healthy tissue to the infarct core (Spearman ρ = -0.92, p = 4.5 × 10^−7^). Tissue-area-corrected analysis confirmed that CNG loss (26% retained in infarct zone vs. normal; p = 0.016) vastly exceeded overall tissue loss, indicating selective vulnerability rather than bulk tissue destruction. Mechanistically, oxidative phosphorylation (OXPHOS) activity in CNG-positive spots correlated positively with CNG survival (partial ρ = 0.65, p = 0.009 after controlling for spot composition), consistent with a high metabolic demand underlying their ischemic sensitivity. Pyroptosis—but not ferroptosis or apoptosis—signatures correlated with CNG loss (partial ρ = -0.67, p = 0.006). These findings identify CNGs as selectively vulnerable cellular components of the cardiac autonomic system in MI and nominate metabolic failure and pyroptosis as candidate mechanisms, providing a spatially resolved foundation for future mechanistic and interventional studies.

## Introduction

Post-MI autonomic remodeling—sympathetic hyperinnervation at border zone, denervation at infarct core—is a well-established driver of arrhythmia and sudden cardiac death. Current research is neuron-centric, focusing on nerve sprouting, growth factors, and axonal regrowth^[1-3]^.

Cardiac nexus glia (CNGs) were identified in 2021 as a population of astroglia-like cells conserved across zebrafish, mouse, and human hearts^[4]^. Originating from the hindbrain neural crest and differentiating via Meteorin–Jak/Stat3 signaling, CNGs express the astroglial markers GFAP, GLAST, and glutamine synthetase, and integrate into the intrinsic cardiac nervous system with net-like morphology apposed to neurons and synapses. Functional studies in zebrafish demonstrated that CNGs regulate heart rate and rhythm through modulation of both sympathetic and parasympathetic branches, and their ablation increases susceptibility to ventricular arrhythmias^[4]^. Beyond development, cardiac glial cells release the neurotrophic factor S100B in response to neuronal damage^[5]^, and glial cells of the sinoatrial node modulate cardiac rhythm in situ^[6]^. Together, these findings establish cardiac glia as functional components of the cardiac nervous system; however, whether CNGs survive or die after myocardial infarction—and how their fate relates to the disrupted autonomic landscape of the infarcted heart—remains entirely unknown.

Here, leveraging publicly available spatial transcriptomic profiling of human infarcted hearts, we mapped CNG distribution across the spatial continuum from normal myocardium to infarct core. We report that CNGs undergo selective, spatially graded loss that exceeds tissue-level destruction, and provide transcriptome-level evidence linking this vulnerability to metabolic (OXPHOS) decline and pyroptosis.

## Results

### 2.1 CNGs are progressively lost from healthy myocardium toward the infarct core

We analyzed 16 human cardiac Visium spatial transcriptomes grouped by sampling location: normal myocardium (Normal, n=4), border zone adjacent to remote myocardium (Outer-BZ, n=3), remote zone (RZ, n=3), infarct zone (IZ, n=5), and border zone adjacent to the infarct (Inner-BZ, n=1) (Fig. 1A). CNG spots were defined as S100B^+^ or GFAP^+^ with undetectable CD68, following established astroglial marker panels for cardiac glia^[4, 5]^. CNG abundance decreased monotonically along the spatial gradient (median CNG percentage: Normal 3.05%, Outer-BZ 2.70%, RZ 1.13%, IZ 0.77%; Fig. 1B). CNG retention correlated strongly with spatial position (Spearman ρ = -0.920, p = 4.47 × 10^−7^; Kruskal-Wallis p = 0.011; IZ vs. Normal, Wilcoxon p = 0.016; Fig. 1B).

**Fig 1.**
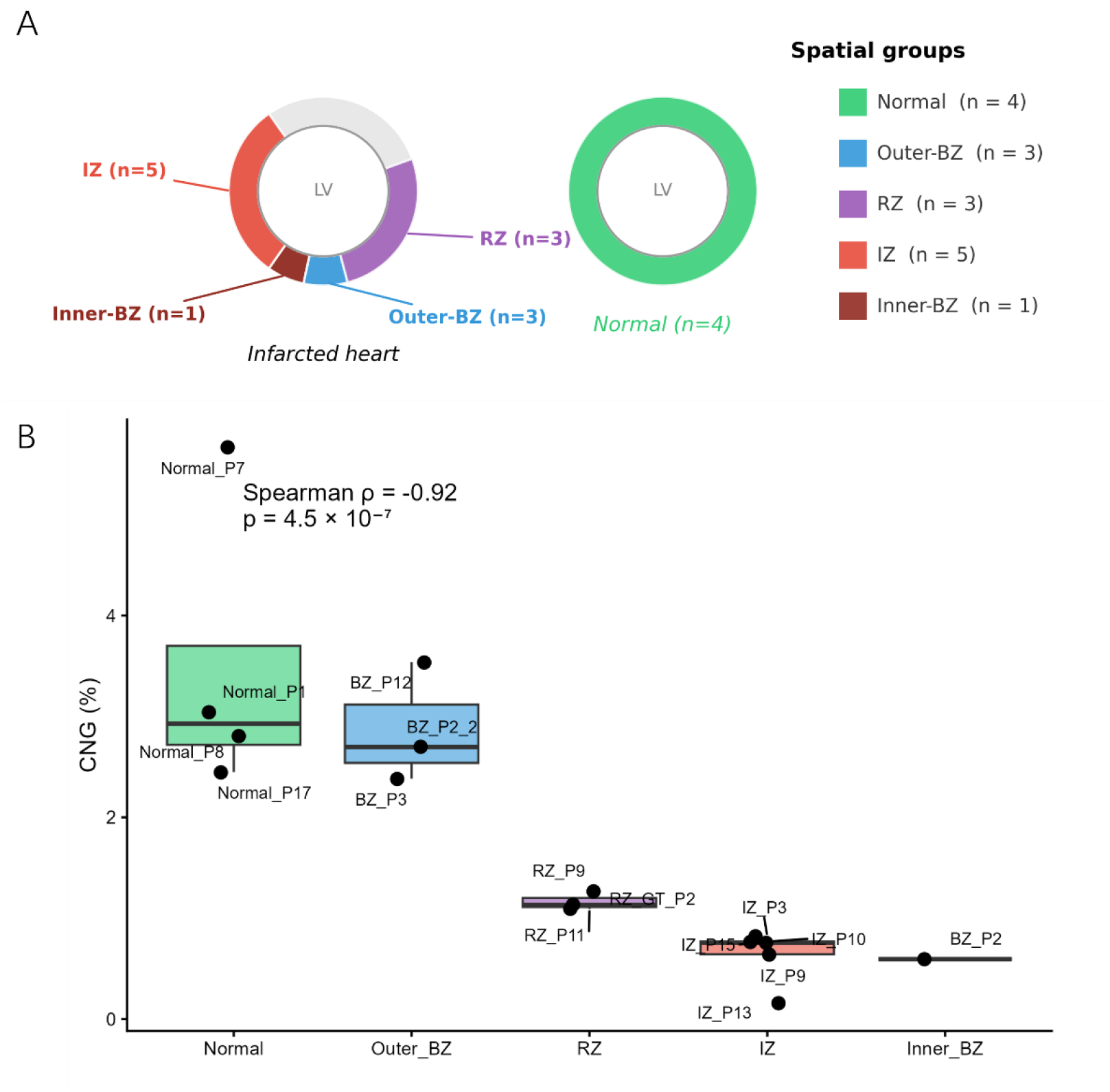
Study design and spatial gradient of CNG loss in human myocardial infarction. (A) Sampling locations and group sizes. Schematic short-axis view of an infarcted heart (not to scale) showing the infarct zone (IZ), border zone adjacent to the infarct (Inner-BZ), border zone adjacent to remote myocardium (Outer-BZ), and remote zone (RZ); right, a normal heart. (B) Percentage of CNG spots (S100B^+^/GFAP^+^, CD68^−^) across the five spatial groups. Boxes, median and interquartile range; points, individual samples (jittered for visibility). CNG percentage declined significantly with proximity to the infarct core (Spearman ρ = -0.92, p = 4.5 × 10^−7^; Kruskal-Wallis p = 0.011).

### 2.5 CNG loss is selective and exceeds overall tissue loss

Total spot counts—a proxy for tissue amount—did not differ between IZ and Normal samples (Wilcoxon p = 0.556), indicating preserved tissue cellularity in our IZ samples. To correct for tissue-area differences, we computed expected CNG numbers per sample (total spots × median Normal CNG density, 2.93%) and expressed observed counts relative to expectation. Even after this correction, CNG retention dropped from 100% (Normal) and 92% (Outer-BZ) to 39% (RZ), 26% (IZ), and 20% (Inner-BZ) (Fig. 2). Thus, CNG loss (∼4-5 fold relative to Normal density) far exceeds bulk tissue loss, revealing a **selective vulnerability of CNGs to infarct injury**.

**Fig 2.**
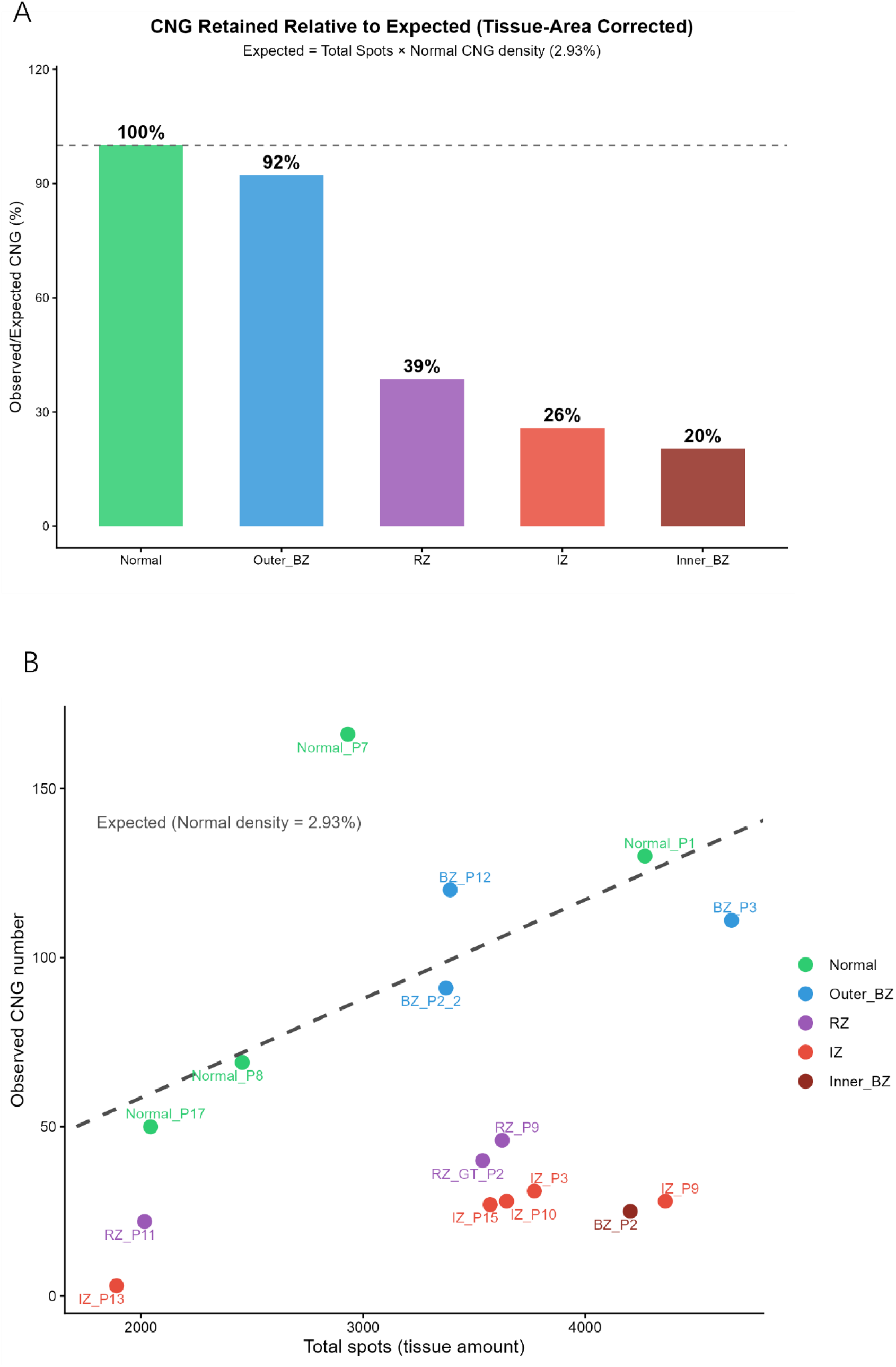
CNG loss is selective and exceeds overall tissue loss. (A) CNG retention (observed/expected) across spatial groups. Expected CNG number per sample = total spots × median CNG density of Normal hearts (2.93%); dashed line, 100% (full retention). Total spot counts (tissue amount) did not differ between groups (IZ vs Normal, Wilcoxon p = 0.556), indicating that declining retention reflects selective CNG loss rather than bulk tissue destruction. (B) Observed CNG number versus total spot number per sample; dashed line, expected CNG number at Normal density. Normal and Outer-BZ samples clustered along the expectation line, whereas RZ, IZ, and Inner-BZ samples fell progressively below it.2.3 | OXPHOS activity and pyroptosis signature associate with CNG survival

### 2.3 OXPHOS activity and pyroptosis signature associate with CNG survival

To explore mechanisms, we scored 15 candidate pathway gene sets within CNG-positive spots and tested their association with CNG retention (Spearman, BH-FDR). OXPHOS showed the strongest positive association (ρ = 0.891, FDR = 5.4 × 10^−5^), while pyroptosis and M2-anti-inflammatory signatures were negatively associated (ρ = -0.694, FDR = 0.014; ρ = -0.703, FDR = 0.014, respectively) (Fig. 3A).

**Fig 3.**
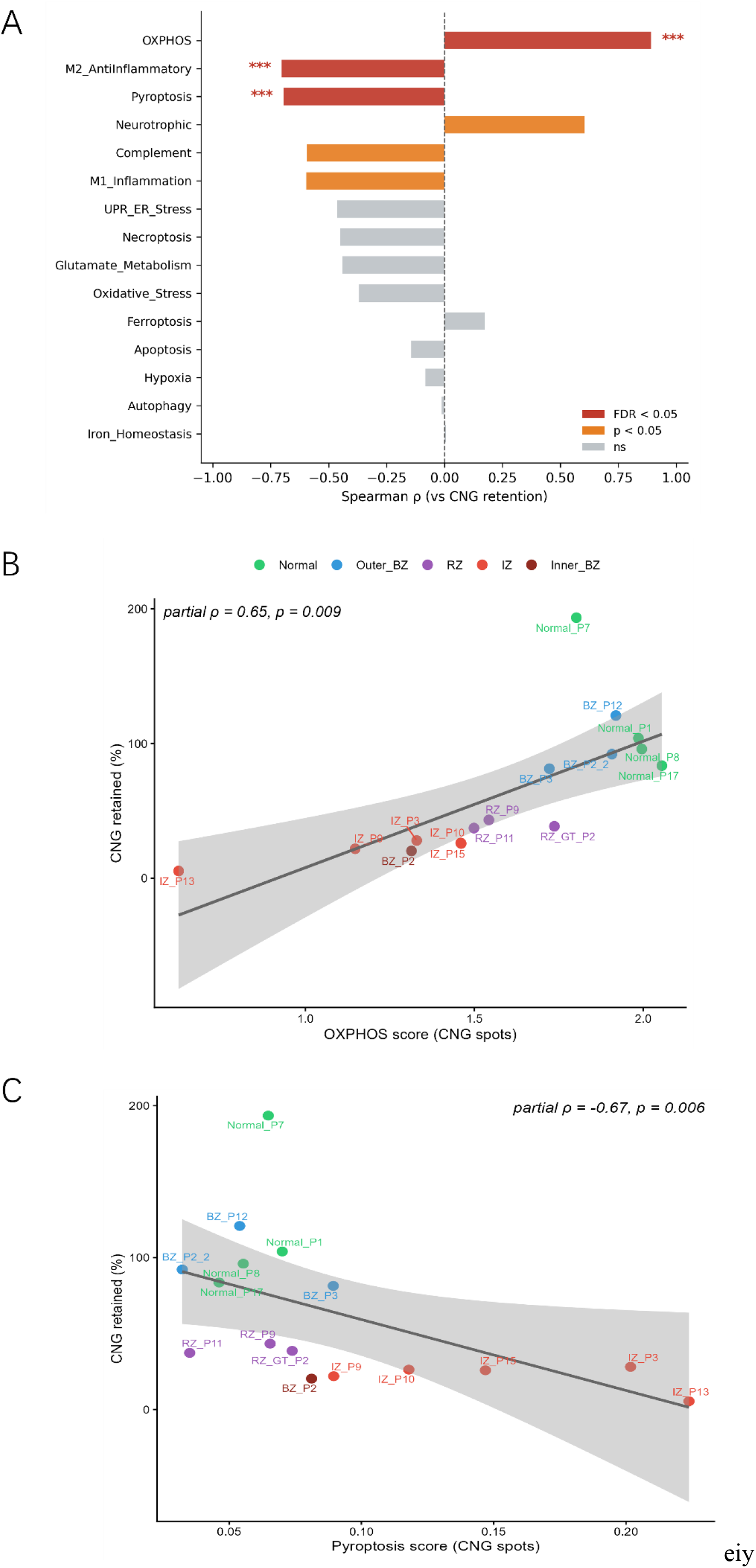
Pathway associations with CNG survival. (A) Spearman ρ of 15 pathway gene sets (scored in CNG-positive spots) versus CNG retention, ordered by effect size; red, FDR < 0.05; orange, p < 0.05; grey, not significant. (B, C) OXPHOS (B) and pyroptosis (C) scores in CNG spots versus CNG retention. Lines, linear fits of raw values (visualization only); reported statistics are partial Spearman correlations controlling for CNG number per sample (OXPHOS: partial ρ = 0.65, p = 0.009; pyroptosis: partial ρ = -0.67, p = 0.006; Bonferroni α = 0.017, 3 tests). The M2 anti-inflammatory signature, despite FDR significance in (A), was abolished after composition control (partial ρ = -0.38, p = 0.156) and is therefore not shown.

Because spot-level measurements mix multiple cell types, decreasing CNG purity in injured samples could inflate immune/inflammation signatures (composition confounding). We therefore recomputed associations controlling for CNG number per sample via partial Spearman correlation (rank-residual method). **OXPHOS remained significantly positive (partial ρ = 0.646, p = 0.009) and pyroptosis significantly negative (partial ρ = -0.671, p = 0.006), both surviving Bonferroni correction (α = 0.017, 3 tests)**, whereas the M2 association disappeared (partial ρ = -0.377, p = 0.156), confirming it as a composition artifact (Fig. 3B,C). In contrast, ferroptosis (ρ = 0.173, p = 0.520), apoptosis (p = 0.594), necroptosis (FDR > 0.05), and hypoxia-response (p = 0.762) gene sets showed no association (Fig. 3A).

These data nominate **metabolic (OXPHOS) failure** as a candidate basis for CNG vulnerability—consistent with the high energetic demands of glial support cells—and **pyroptosis** as the associated death modality, while arguing against ferroptosis at the transcriptomic level.

## Discussion

We present the first spatially resolved map of CNG fate in human myocardial infarction. Three findings stand out: (i) CNG loss follows a steep spatial gradient toward the infarct core; (ii) the loss is selective, exceeding tissue-level destruction by ∼4-5 fold; (iii) it tracks with OXPHOS decline and a pyroptosis signature independently of spot composition.

The high-OXPHOS phenotype of surviving CNGs suggests these cells are metabolically demanding support cells, akin to brain astrocytes (which share GFAP/S100B/GLUL expression and depend on oxidative metabolism). Ischemia-driven metabolic failure could therefore render CNGs disproportionately vulnerable—a testable explanation for their selective loss. The independence of the pyroptosis association from composition confounding nominates inflammasome-mediated death as a candidate execution mechanism; notably, pyroptosis in cardiomyocytes after MI is well documented^[7, 8]^, but its occurrence in cardiac glia has not been examined.

Conceptually, our findings shift attention from neurons to glia in post-MI autonomic remodeling. Given that CNG loss in model organisms increases arrhythmia susceptibility, CNG depletion in human MI may represent an under-recognized contributor to autonomic imbalance and sudden cardiac death, and a candidate therapeutic target.

Several limitations should be noted. First, no markers currently distinguish intracardiac glial cells from other peripheral glia^[5]^. Our CNG definition therefore relied on the S100B/GFAP astroglial panel combined with CD68 exclusion; SOX10, the lineage-defining marker, was below detection in spatial data. Protein-level confirmation by S100B/SOX10 double immunostaining is required and planned. Second, Visium spot mixing limits cell-attribution; we mitigated this via partial correlation, but single-cell/spatial proteomics are needed. Third, the Inner-BZ group comprised a single sample. Fourth, all associations are correlative; causal tests of metabolic failure and pyroptosis in CNG biology await experimental manipulation.

Future work should establish CNG death modalities at the protein level, define the causal contribution of CNG loss to post-MI autonomic remodeling and arrhythmia, and explore protective strategies.

## Methods

### Data collection

We obtained publicly available human cardiac Visium spatial transcriptome datasets profiling myocardial infarction and control hearts^[4]^. Eighteen samples were analyzed. Two were excluded from downstream analysis: one with a corrupted file (failed data import), and one remote-zone sample (RZ_P6) whose CNG percentage (4.42%) exceeded four-fold the dataset maximum with ambiguous sampling annotation in the source metadata. The remaining 16 samples were grouped by documented sampling location: normal myocardium (n = 4), border zone adjacent to remote myocardium (Outer-BZ, n = 3), remote zone (RZ, n = 3), infarct zone (IZ, n = 5), and border zone adjacent to the infarct (Inner-BZ, n = 1).

### CNG identification and quantification

Spatial transcriptomes were processed with Seurat (v5.5.1): normalization by LogNormalize (default parameters). Following established astroglial marker panels for cardiac glia (GFAP, GLAST, glutamine synthetase;)^[9]^ and S100B as a cardiac glial marker^[5]^, we defined CNG spots as those with S100B or GFAP expression detected (> 0) and undetectable CD68. SOX10, the neural crest lineage marker, was below detection sensitivity in spatial data and could not be used. Per sample, we recorded CNG spot counts, total spot numbers, and CNG percentage.

### Tissue-area correction

To test whether CNG loss merely reflects bulk tissue destruction, we computed expected CNG numbers per sample as total spots × median CNG percentage of Normal samples (2.93%), and expressed observed counts as retention = observed / expected × 100%. Total spot counts across groups were compared by Wilcoxon rank-sum test (exact).

### Gene set scoring

Fifteen candidate pathway gene sets were curated (Supplementary Table S1). For each set, genes detectable in a given sample (≥ 3) were averaged (mean of LogNormalized expression) within CNG spots to yield a per-sample score.

### Correlation and confound control

Associations between gene set scores and CNG retention were assessed by Spearman rank correlation with Benjamini-Hochberg FDR correction across 15 sets. To control for spot composition (fewer CNG cells per spot in injured samples inflating non-CNG signals), we computed partial Spearman correlations controlling for per-sample CNG number by the rank-residual method: variables were rank-transformed, x and y were regressed on the control z, and Pearson correlation of the residuals was tested (t = r√(n−3)/√(1−r^2^), df = n−3). Gene sets passing FDR then underwent partial correlation with Bonferroni correction (α = 0.017, 3 tests).

### Additional statistics

Group comparisons by Kruskal-Wallis test; pairwise by two-sided Wilcoxon rank-sum test. Gradient trends by Spearman correlation with ordered spatial position. α = 0.05 throughout.

## Supporting information

Supplementary Table S1. Curated pathway gene sets used in this study.

## Software and data availability

Analysis was performed in R 4.6.0 (Seurat 5.5.1, anndata, ggplot2). Processed data tables and analysis code are deposited at Zenodo (https://doi.org/10.5281/zenodo.22759503)

